# Academic stress through salivary biomarkers: A multivariate analysis of cortisol, IL-1β, CRP, and IgA levels and their variations as a function of sex

**DOI:** 10.1101/2024.10.16.618261

**Authors:** Rodrigo Castillo Klagges, Camila Pezo Sáez, Luis Aguila, Verónica Pantoja, Favián Treulen Seguel

**Affiliations:** Escuela de Tecnología Médica, Facultad de Medicina y Ciencias de la Salud, Universidad Mayor, Temuco-Chile; Centro de Excelencia de Biotecnología en Reproducción, Facultad de Medicina, Universidad de La Frontera, Temuco-Chile; Magíster en Neurociencias de la Educación, Escuela de Educación, Facultad de Ciencias Sociales y Artes, Universidad Mayor, Temuco-Chile

**Author notes:** Autor de correspondencia: Favián Treulen, Escuela de Tecnología Médica, Facultad de Medicina y Ciencias de la Salud, Universidad Mayor, Temuco-Chile.

**Keywords:** Academic stress, cytokines, C-reactive protein, immunoglobulin A, inflammation, interleukin 1-beta, salivary biomarkers, SISCO

## Abstract

**Introduction:** Academic stress can activate physiological changes mediated by the sympathetic nervous system (SNS) and the hypothalamic-pituitary-adrenal (HPA) axis, triggering the release of biomarkers such as cortisol and proinflammatory cytokines. Although physiological stress has been studied in relation to different inducers and diseases, there is still a gap regarding the association of academic stress with biological markers. Thus, this study aimed to associate the levels of academic stress against biological markers isolated from saliva from undergraduates’ students.

**Materials and methods:** 81 students (53 women and 28 men) were recruited and completed the SISCO inventory to determine the level of academic stress. The levels of cortisol, interleukin-1β (IL-1β), C-reactive protein (CRP) and immunoglobulin A (IgA) from saliva samples were determined by ELISA assays, and data were analyzed using ANOVA, Pearson correlation tests. A predictor model was estimated by lineal regression.

**Results:** Stress categorization following the SISCO inventory showed that 37% of the students grouped in the low stress level (<48%), 35% grouped in the moderate stress level (>49% <60%), and 28% in high stress level (>61% <100%). The levels of salivary markers were similar across stress categories, however the trends identified—such as the decrease in cortisol and the increase in pro-inflammatory markers in male participants categorized in the high stress group—suggest a possible association between these biomarkers with academic stress gender-dependent. The multivariable model including the 4 biomarkers resulted in R^2^ = 0.14 with predictions that were roughly within +/-20% of stress levels.

**Conclusion:** In conclusion, no significance was found in the association of salivary biomarkers with academic stress levels. However, trends were observed with increasing levels of academic stress in men. The concentration of these biomarkers may be affected by sex. Further research will consider individual factors, longitudinal assessments, and the use of multiple psychometric tools to better define the interaction between academic stress and salivary biomarkers.

## 1. Introduction

“A stressor event” could be considered as any stimulus that the brain interprets as a threat or challenge (1). The compensatory reaction to this stimulus is called “stress response,” which is necessary for the survival of species, it prepares the organism for physical or psychological events, such as fear, tension, or danger situations (2). This response is an adaptative change that involves the activation of the sympathetic nervous system (SNS) and the hypothalamic-pituitary-adrenal (HPA) axis (3).

The activation of the SNS causes a rapid short-term physiological modulation, increasing glycemia, blood pressure, heart rate, and stimulating the inflammatory and immune response (4–6). This initial response is regulated by the HPA axis and its final hormone, cortisol, which has an anti-stress and anti-inflammatory function (6,7). Cortisol plays a fundamental role in the body’s response to stress (8). Due to its ability to inhibit leukocyte proliferation, pro-inflammatory proteins, and antigen presentation (9–11). Similarly, acute stress also activates the immune system (12), raising immunoglobulin A (IgA) in saliva as a countermeasure against potential exposure to pathogens in a “fight or flight” scenario (6,13).

Stressful events are common in daily life. However, humans can control what we perceive as stressful and how we respond to it (14,15). Exaggerated or recurring negative interpretations such as worry, magnification, and helplessness are maladaptive catastrophic responses to stress that can prolong cortisol secretion (16,17). This chronic activation and the repeated waves of the HPA axis cause hypercortisolism (18), generating resistance in glucocorticoid receptors and increasing their affinity for mineralocorticoid receptors, triggering pro-inflammatory effects (1,19–21).

These physiological markers can be determined in blood samples, evaluating the levels of pro-inflammatory cytokines (e.g., IL-1β) and acute phase proteins (e.g., C-reactive protein [CRP]). In addition, these inflammatory markers are also detected in saliva in response to acute and chronic psychological stress (22–27). Moreover, the physiological effects of stress, such as the already mentioned link between chronic stress and a systemic pro-inflammatory state (28), is considered a risk factor for diverse pathologies such as cardiovascular disease, hypertension, diabetes, and cancer (29-33).

In the biomedical context, “academic stress” is considered as a set of daily stressors events that influence the psychological and physiological well-being of students. From this perspective, academic milestones such as exams, tests, and oral presentations are the most relevant triggers during student life, causing significant behavioral, cognitive, and physiological-emotional consequences, contributing to chronic stress state and mental health deterioration (34–37). Additionally, there are many other stress factors that can affect student performance, such as academic demands or family environment among others (38). It is known that stressful situations in students trigger psychological and physical symptoms such as anxiety, fatigue, insomnia, and signs associated with academic performance (39). Although the detection of academic stress is particularly complex, as psychological, social, and biological factors are involved (40), tools focusing on the stressor potential of different academic conditions have been developed, such as the SISCO inventory of academic stress (SISCO-AS) (41), which was updated in 2018 to its second version (SISCO-II-AS) (42). This tool has been used in several studies in South America, demonstrating valid and reliable results (43-45).

For all the aforementioned reasons, it is essential to understand the physiological effects of academic stress in order to design detection methods and preventive interventions to mitigate the harmful effects. Although some studies have shown that salivary biomarkers are modified in individuals under acute and/or chronic stress, it has not been directly linked to academic stress. The objective of this research is to evaluate the association of salivary biomarkers with levels of academic stress in undergraduates’ students.

## 2. Materials and methods

### 2.1. Sample and participants

The participant cohort were student volunteers from the Faculty of Medicine and Health Sciences at Universidad Mayor, Temuco, Chile. A group of eighty-one students, aged between 18 and 30 years, was selected, including 53 women and 28 men. The following exclusion criteria were considered to avoid directly or indirectly affecting the systemic inflammatory state (46-48): pregnancy and/or breastfeeding, acute and/or chronic infections, chronic diseases, endocrine diseases, smoking, drug and/or alcohol abuse, and the use of any type of medication. Participants were invited to voluntarily participate by means of posters displayed at the university with a QR code, which redirected them to an online survey. This survey allowed the researchers to evaluate the inclusion/exclusion criteria of the participants and to collect personal information such as name, gender, age, career, current academic year, telephone number and e-mail. The selected participants were contacted 3 days prior to sample collection and instructed to abstain from smoking, alcohol, and exercise. To avoid saliva contaminated with blood or other interferents, participants were instructed not to eat, drink, or brush their teeth for a period of two hours prior to sample collection.

### 2.2. Study design

The participants were recruited from Monday, October 30 to November 9 (Monday-Friday), from 9:30 am to 13:00 pm. This period corresponds to the end of the academic semester. They were asked to sign an informed consent form, which was reviewed and approved by the Scientific Ethical Committee of the Universidad Mayor (Resolution No. 0398). Then, the SISCO inventory was applied online. Participants were instructed to rinse their mouth with water, 10 minutes before sampling, and 1.5 ml of salivary sample was collected in a 2 mL microcentrifuge tube. The collected samples were identified with the participant’s order number and the date of extraction, centrifuged at 1000 g for 10 minutes to remove cell debris, mucin, and debris. Subsequently, they were stored at - 20ºC until analysis.

#### 2.2.1 SISCO

Participants were administered the SISCO-II-AS survey with 33 items that inquired about the three systemic-processual components of academic stress: stressors, symptoms and coping strategies. The first item, in dichotomous terms (yes-no), allows to determine whether the respondent is a candidate or not to answer the inventory. The second item made it possible to identify the level of intensity of academic stress. Eight items, Likert-type scale of six categorical values, allow identifying the frequency in which the demands of the environment are valued as stressful stimuli. Another 17 items allow to identify the frequency with which symptoms or reactions to a stressor stimulus occur.

Finally, six items allow to identify the frequency of use of coping strategies. The last three sections use a Likert scale (1: never, 5: always). This survey made it possible to classify the participants into the following groups: low (<48%), moderate (>49%<60%) and high academic stress level (61%><100%). The sections of this instrument can be used as a whole, separately or combined (49).

#### 2.2.2 Biomarker analysis and reagents used

The following stress biomarkers were analyzed: Cortisol (Cortisol ELISA Kit Abcam, Waltham, MA, United States), IL-1β (Human IL-1 beta ELISA Kit Abcam, Waltham, MA, United States), PCR (Human C Reactive Protein ELISA Kit Abcam, Waltham, MA, United States), IgA (Human IgA ELISA Kit Abcam, Waltham, MA, United States). A microplate reader (BioTek Instruments EL800, Winooski, VT, United States) and its integrated analysis software (BioTek Gen5 software, Winooski, VT, United States) were used for reading results. The determinations of each biomarker were performed according to the manufacturer’s recommendations.

### 2.3 Statistical analysis

The statistical program GraphPad Prism for Windows (Version 10, GraphPad Software, Boston, USA) was used to analyze the results. The normality of the data obtained was evaluated using Bartlett’s test and the Brown-Forsythe test. To evaluate the relationship between variables, Pearson correlation coefficient was calculated to assess the relationship between markers and academic stress percentage. Simple linear regression analysis model was used to assess the correlation markers and academic stress percentage. To compare differences between stress groups, one-way ANOVA and a multiple comparison test were applied. Unpaired *t* test was used to perform the comparison between biomarkers levels and sex. *P*<0.05 was considered statistically significant. The results are expressed as mean ± SD.

Individual predictors were determined using analysis of variance (using the lm()-function in R, Vienna, Austria), and the coefficients of determination were used for numerical comparison of the variables. Finally, a multivariable model was constructed including all 4 variables.

## 3. Results

Descriptive statistics of participants according to sex, career and academic stress levels are shown in Table 1. Briefly, from the eighty-one students surveyed, presented a gender distribution of whom 65.4% (53/81) women and 34.6% (28/81) men (Table 1). The distribution by academic stage (seen as cycle/year of completion) was as follows from the first cycle/year until to fifth or last cycle/year: 25.9%, 23.5%, 23.5%, 23.5%, and 3.6%.

**Table 1.** Distribution of participants according to sex, career and academic stress mean percentage.

| Career | Men | Women | AS-MP |
| --- | --- | --- | --- |
| Bachelor's degree in health sciences | 1 | 2 | 55% |
| Nursing | 4 | 10 | 59% |
| Phonoaudiology | 2 | 3 | 63% |
| Kinesiology | 1 | 5 | 61% |
| Medicine | 9 | 4 | 60% |
| Nutrition and Dietetics | 0 | 3 | 67% |
| Obstetrics | 0 | 13 | 68% |
| Dentistry | 2 | 1 | 64% |
| Medical Technology | 9 | 9 | 64% |
| Occupational therapy | 0 | 3 | 70% |
| **Total** | **28** | **53** | **63%** |
|  | **81** |  |  |

AS-MP Academic stress mean percentage

The SISCO survey was used as a tool to assess the level of academic stress, focusing on the frequency of stressors and associated symptoms. This tool showed that 76.8% of the surveyed answered that: any type of evaluation is “always” or “almost always” a disturbing factor. In addition, 47.3% stated that they felt uneasy “almost always” due to task overload.

As shown in **Table 1**, the results showed that most degrees associated with healthcare present an elevated level of academic stress. Obstetrics, Occupational therapy, and Human Nutrition were the careers with the highest percentage of academic stress. Regarding the stress level classification, 37% of the participants showed a low level of stress, with an average stress level of 41.2%. Similarly, 34.6% presented a moderate level of stress, with a mean stress of 55%. On the other hand, 28.4% showed a high level of academic stress, with a mean stress of 68.2%. The gender distribution according to the levels of academic stress showed that 63.3% at the low level were women, at the moderate level 60.7% were women and 39.3% were men. Finally, at the high level 73.9% were women and 26.1% were men.

When analyzing biomarker levels (cortisol, IL-1β, CRP, and IgA) across academic stress categories (low, medium, and high levels) (**Figure 1**), there was an evident decrease in cortisol and a tendency of IL-1β and CRP to increase in males as the level of academic stress rises. However, no significant difference was observed.

**Figure 1.**
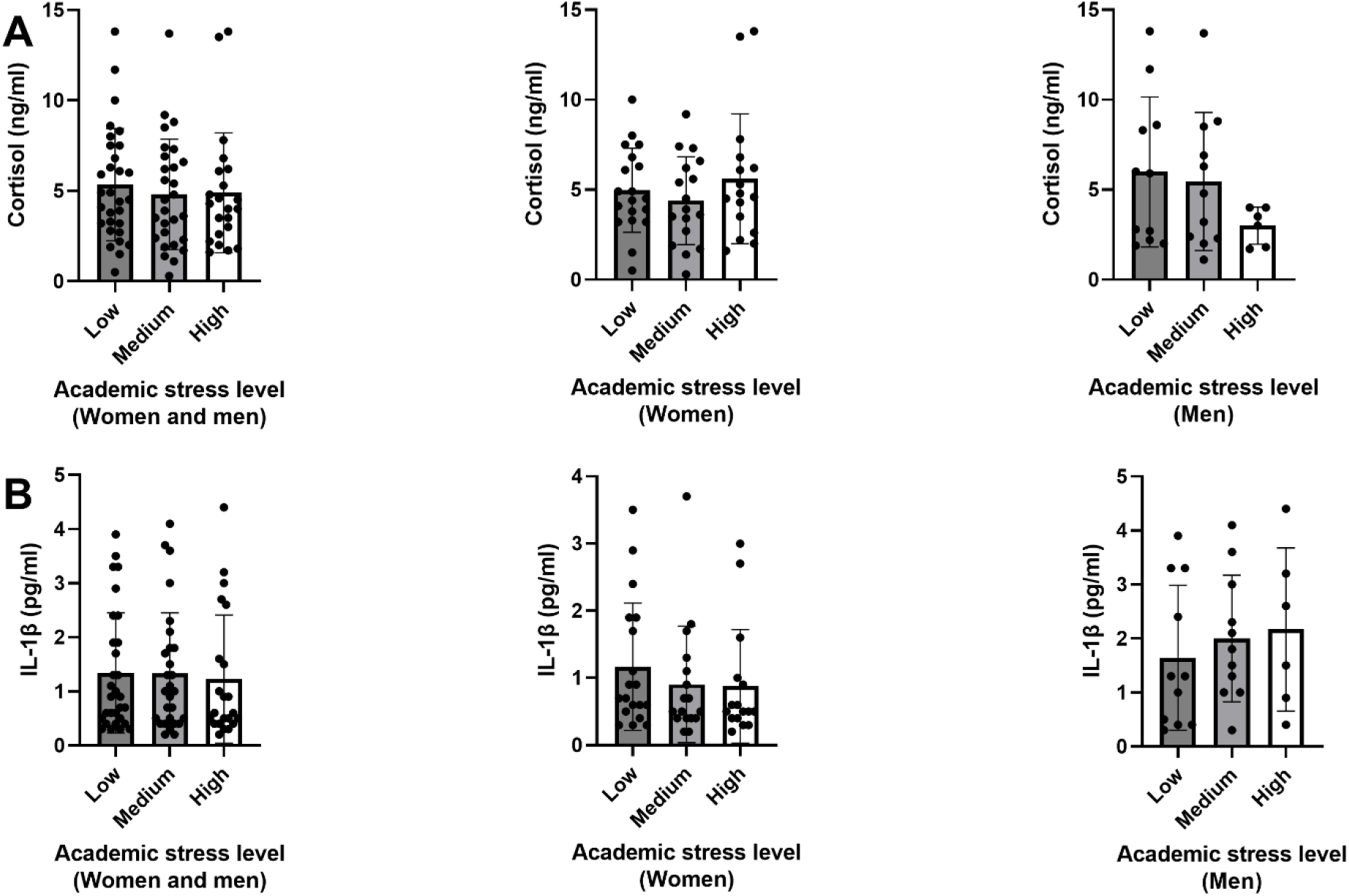
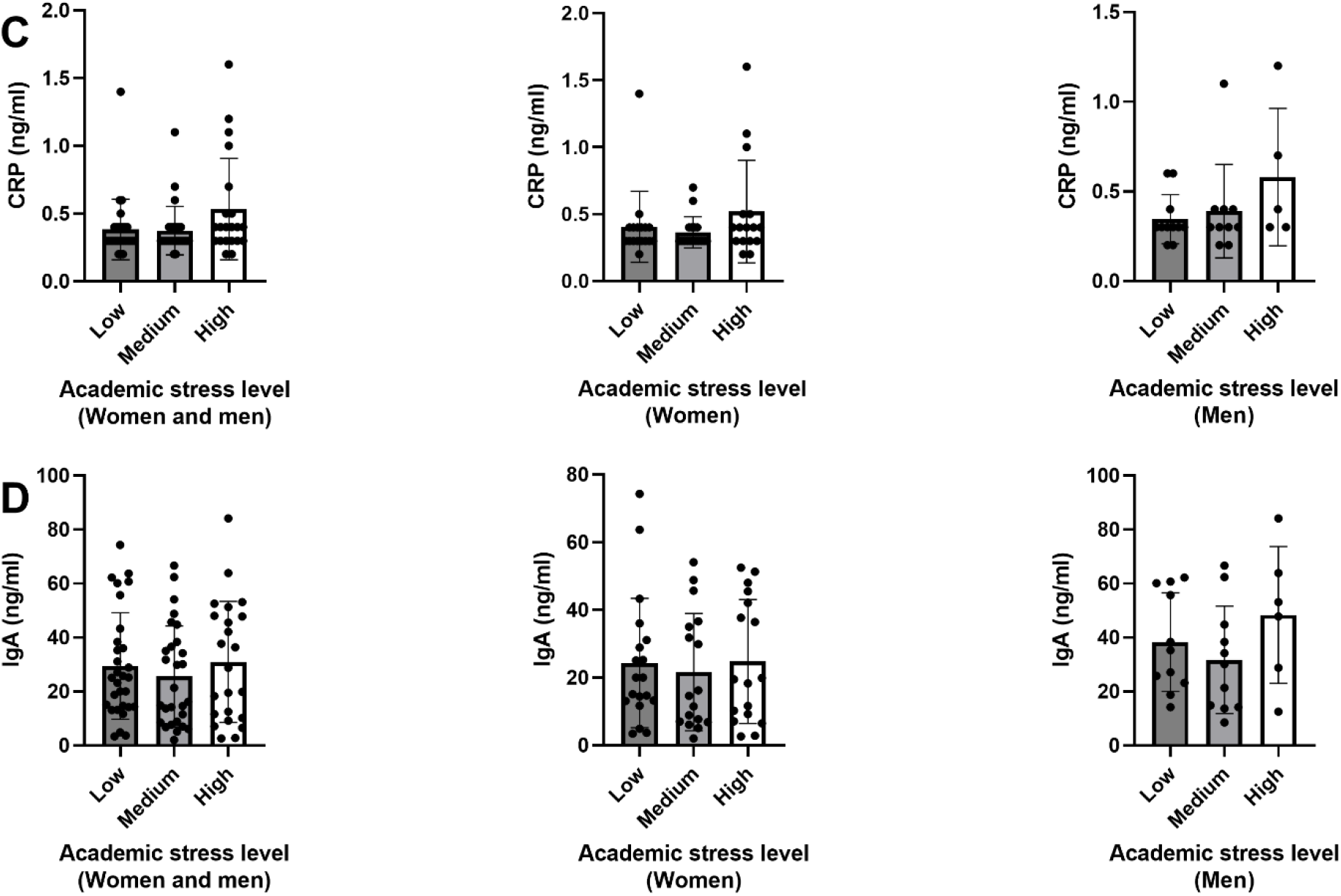
Bar plots illustrate the analysis of stress markers in saliva across levels of academic stress determined according to the SISCO inventory. Cortisol (A), IL-1β (B), CRP (C) and IgA (D). Mixed and separated by sex. Results are expressed as mean ± SD.

A simple linear regression model was used in normally distributed data to determine the relationship between each biomarker and the percentage of academic stress. (**Figure 2**). Cortisol in males showed a moderate negative correlation with academic stress level percentage (*P*=0.28). In contrast, IL-1β and IgA in males showed a positive correlation with academic stress level percentage (*P*=0.57 and *P*=0.46, respectively). Likewise, in females, a negative trend was observed in the correlation of IL-1β. Interestingly, it was observed that in the comparison of the concentrations of the studied biomarkers based on gender (**Figure 3**), it revealed that men presented significantly higher levels of IL-1β and IgA in saliva than women.

**Figure 2.**
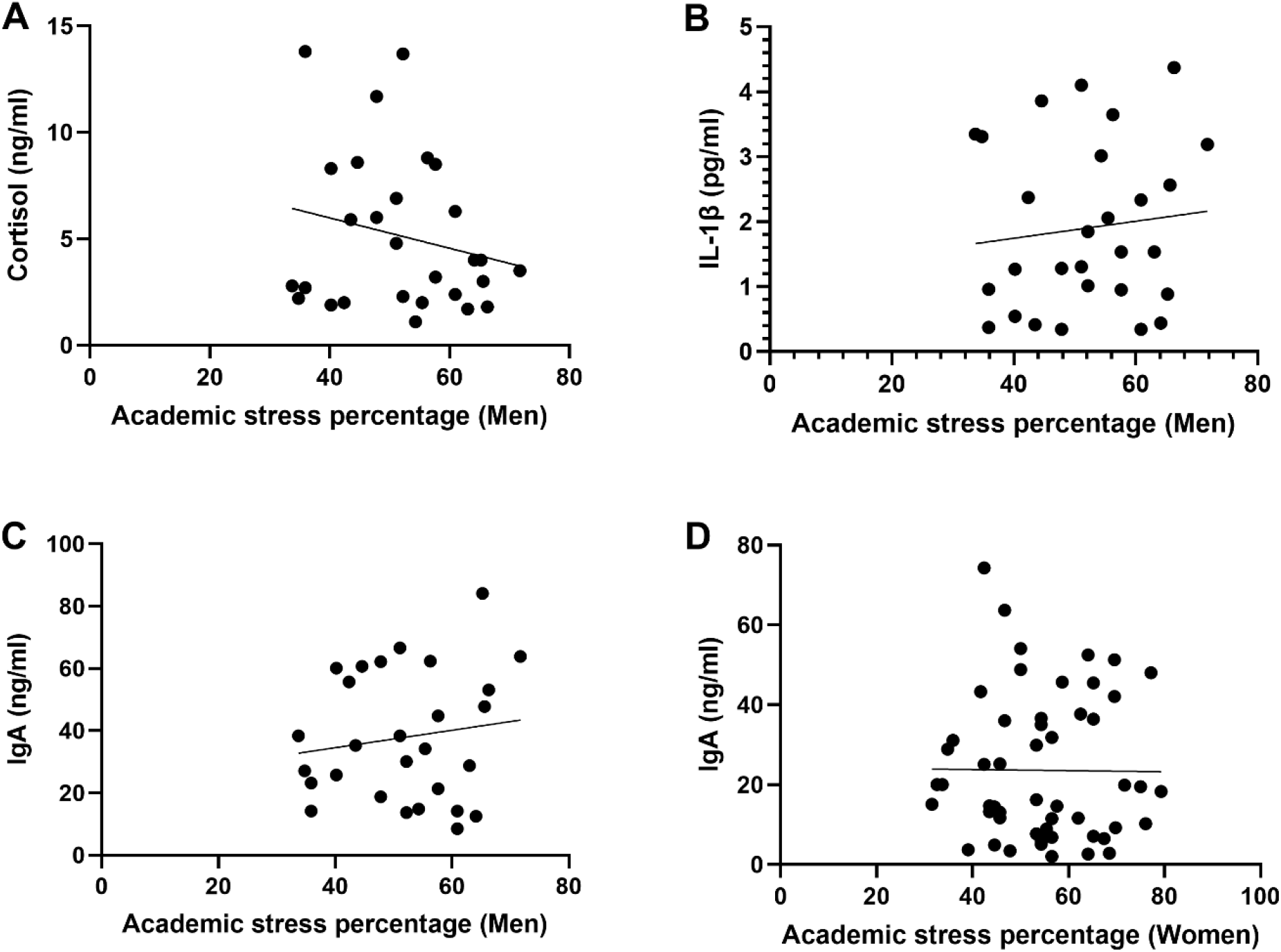
The scattergrams of the linear regression illustrating the distribution and relationship between the concentration of cortisol in men (A), IL-1β in men (B), IgA in men (C) and IgA in women (D), and the academic stress level percentage. Results are expressed as mean ± SD. The significance level was set for a *P*<0.05.

**Figure 3.**
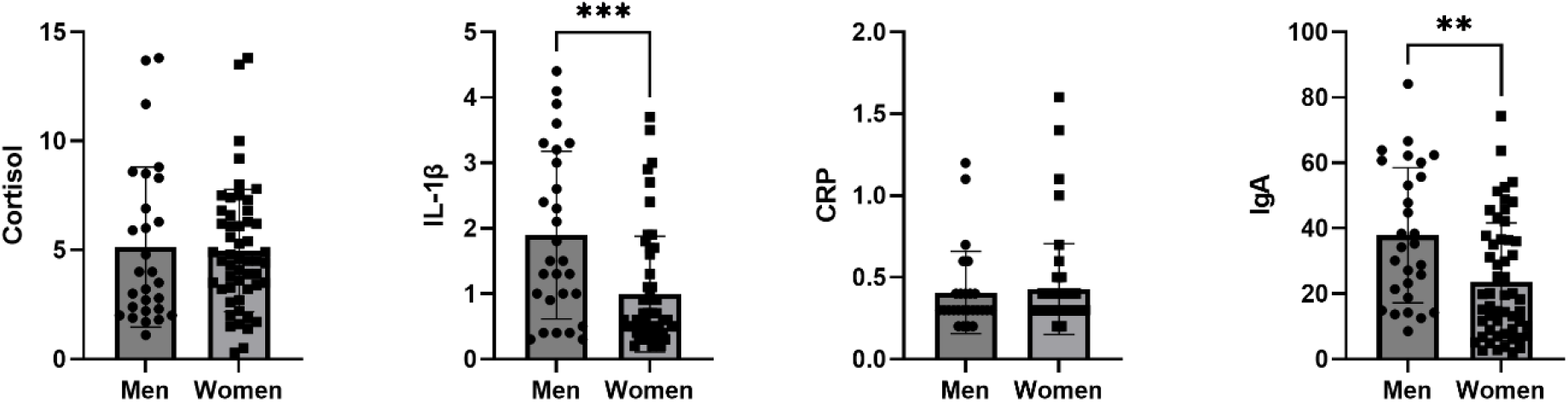
T test plot, comparing the mean concentration of cortisol (A), IL-1β (B), CRP (C) and IgA (D), in men and women. Results are expressed as mean ± SD. The significance level was set for a *P*<0.05. *** (*P*=0.0002), ** (*P*=0.0018).

Then, a multivariable predictor model including stress biomarkers measurements was constructed thus:

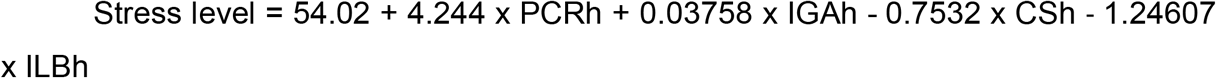

The multivariable model had an R squared= 0.14 with predictions ranging approximately within +/- 20% of the stress levels.

## 4. Discussion

In recent years, the use of salivary biomarkers to determine biological stress has been widely supported in the literature for its advantages over other types of biological samples (50-52), offering a noninvasive and accessible way to measure the physiological response to stress. This study aimed to identify correlations between biological markers of stress (cortisol, interleukin 1-beta, C-reactive protein, and immunoglobulin A) detected in saliva and academic stress determined by the SISCO survey in undergraduate students.

Although, no statistical correlation was found between the levels of salivary markers and stress, it was seen a trend between the level of stress levels and concentrations of cortisol, IL-1β and CRP in males. The absence of statistical strength could be affected by different factors, such as variability in the stress response among individuals and/or the psychometric test applied. This does not rule out the relevance of these biomarkers in academic stress contexts, because several studies have shown that these biomarkers are modified in response to acute or chronic psychological stress (7,26,53,54).

Salivary cortisol, as a reliable indicator of stress, is closely correlated with plasma concentrations (55). Interestingly, salivary cortisol tended to decrease in the high academic stress group. The concentration of cortisol increases in saliva after acute stress (44), however, in chronic periods or during stages of constant secretion, the cortisol levels decrease due to desensitization of the HPA axis and the presence of a proinflammatory state (56,57). This phenomenon of HPA-axis depletion or glucocorticoid receptor resistance could explain the above state downward trend. Similarly, another study observed that men trigger a greater response to stress than women (58), as well as morning salivary cortisol is a predictor of academic stress in male undergraduates (59). On the other hand, it has also been indicated that women tend to report greater subjective stress (60), we believe that this could be a factor which affected the objectivity of the results obtained from the SISCO survey in women (61).

As cortisol, the inflammatory markers interleukin-1 beta and C-reactive protein have been used to assess inflammatory response to stress. Considering the upward trend of these biomarkers and the downward trend of cortisol, it could reflect the same dynamics pointing to prolonged activation of the HPA axis and the SNS (61). It is also described that men tend to show more pronounced inflammatory responses to stress compared to women, which is consistent with the higher levels of IL-1β and IgA in men than in women recorded here. This aspect should be explored in future studies to better understand the mechanisms underlying these sex differences. On the other hand, studies have observed an increase in IL-1β after stress exposure, showing that this cytokine has a high sensitivity for the detection of acute stress (21,26,62). Similarly, salivary levels of IL-1β were correlated with insomnia in undergraduates (63). Same way, another study using chronically stressed animals reported that IL-1β mRNA expression from the submandibular glands was closely correlated with chronic stress (64). Moreover, a study in men demonstrated a significant increase in CRP concentration in response to a stressful task, suggesting an important role in dysregulation caused by stress (65). However, further studies are needed to determine whether there is a relationship between IL-1β and CRP concentrations with academic stress levels. This demonstrates that the measurement of academic stress is complex and cannot be categorized simply as acute or chronic stress.

Salivary immunoglobulin A (sIgA) is another potential biomarker of great relevance, playing a crucial role in the immune system, particularly in mucosal immunity. Studies show that sIgA increases in response to acute stress situations (66,67), this response may be particularly useful in studies of academic stress, given that students frequently face situations that trigger acute stress responses, such as exams and oral exposures (68). Similarly, another study found that the level of sIgA changes in response to psychological factors, decreasing in subjects with elevated levels of perceived stress (7).

In addition, we present a validated estimator for academic stress. The multivariable model responds in 14% of the change of academic stress. Other studies in college students have established predictive models of post-traumatic stress with correlations ranging from 19 to 38%. (69,70). Thus, aunque el valor de R del presente estudio es relativamente bajo, nuestro estudio permite establecer un modelo de regresión del estrés académico en torno a marcadores biológicas. However, the strength of the proposed model must be increased.

Finally, a limitation of our study lies in a single time point sampling and the number of samples (insufficient quantity of biological replicates). Although these clear limitations, this is a pioneer study generating a statistical predictor of academic stress by measuring salivary markers. Therefore, our findings should be mainly taken in the context of academic stress. Second, the SISCO survey comprised only academic stressors and stress reactions, but psychological stress is a multi-factorial response (71-73).

## 4. Conclusions

In conclusion, although all academic stress categories (low, medium and high) showed similar levels of the salivary biomarkers assessed, it is possible to establish a predictor model correlating salivary biomarker levels with sex-dependent academic stress. Further studies will be focused on increasing study population, considering multiple assessment tools and/or lineal mixed models to elucidate interactions between academic stress and students’ physiological responses.

## Supporting information

Suplemental file correlation CRP IL-1B men

## References

1. Hannibal KE, Bishop MD. Chronic Stress, Cortisol Dysfunction, and Pain: A Psychoneuroendocrine Rationale for Stress Management in Pain Rehabilitation. Phys Ther. 2014 Dec 1;94(12):1816–25.

2. Torrades S. Estrés y burn out. Definición y prevención. Offarm [Internet]. 2007 Nov 1 [cited 2022 Nov 10];26(10):104–7. Available from: http://www.elsevier.es/es-revista-offarm-4-articulo-estres-burn-out-definicion-prevencion-13112896

3. Russell G, Lightman S. The human stress response. Nat Rev Endocrinol. 2019 Sep 27;15(9):525–34.

4. Vinik AI, Maser RE, Ziegler D. Autonomic imbalance: prophet of doom or scope for hope? Diabetic Medicine. 2011 Jun 16;28(6):643–51.

5. Godoy LD, Rossignoli MT, Delfino-Pereira P, Garcia-Cairasco N, de Lima Umeoka EH. A Comprehensive Overview on Stress Neurobiology: Basic Concepts and Clinical Implications. Front Behav Neurosci. 2018 Jul 3;12.

6. Lambert M, Couture-Lalande MÈ, Brennan K, Basic A, Lebel S, Bielajew C. Salivary secretory immunoglobulin A reactivity: a comparison to cortisol and -amylase patterns in the same breast cancer survivors. Współczesna Onkologia. 2018;22(3):191–201.

7. Engeland CG, Hugo FN, Hilgert JB, Nascimento GG, Junges R, Lim HJ, et al. Psychological distress and salivary secretory immunity. Brain Behav Immun. 2016 Feb;52:11–7.

8. Dhama K, Latheef SK, Dadar M, Samad HA, Munjal A, Khandia R, et al. Biomarkers in Stress Related Diseases/Disorders: Diagnostic, Prognostic, and Therapeutic Values. Front Mol Biosci. 2019 Oct 18;6.

9. Rhen T, Cidlowski JA. Antiinflammatory Action of Glucocorticoids — New Mechanisms for Old Drugs. New England Journal of Medicine. 2005 Oct 20;353(16):1711–23.

10. Strehl C, Ehlers L, Gaber T, Buttgereit F. Glucocorticoids—All-Rounders Tackling the Versatile Players of the Immune System. Front Immunol. 2019 Jul 24;10.

11. Dra. Miriam Sánchez Segura LicRMGGDraVMS y DraCMA. Asociación entre el estrés y las enfermedades infecciosas, autoinmunes, neoplásicas y cardiovasculares. Revista Cubana de Hematología, Inmunología y Hemoterapia. 2006;22:1561–2996.

12. Castro-Quintas Á, Palma-Gudiel H, San Martín-González N, Caso JR, Leza JC, Fañanás L. Salivary secretory immunoglobulin A as a potential biomarker of psychosocial stress response during the first stages of life: A systematic review. Front Neuroendocrinol. 2023 Oct;71:101083.

13. Giacomello G, Scholten A, Parr MK. Current methods for stress marker detection in saliva. J Pharm Biomed Anal. 2020 Nov;191:113604.

14. Hirsch CD, Barlem ELD, Almeida LK de, Tomaschewski-Barlem JG, Lunardi VL, Ramos AM. FATORES PERCEBIDOS PELOS ACADÊMICOS DE ENFERMAGEM COMO DESENCADEADORES DO ESTRESSE NO AMBIENTE FORMATIVO. Texto & Contexto - Enfermagem. 2018 Mar 5;27(1).

15. Denson TF, Spanovic M, Miller N. Cognitive appraisals and emotions predict cortisol and immune responses: A meta-analysis of acute laboratory social stressors and emotion inductions. Psychol Bull. 2009 Nov;135(6):823–53.

16. Johansson A, Gunnarsson L, Linton SJ, Bergkvist L, Stridsberg M, Nilsson O, et al. Pain, disability and coping reflected in the diurnal cortisol variability in patients scheduled for lumbar disc surgery. European Journal of Pain. 2008 Jul 9;12(5):633–40.

17. Müller MJ. Helplessness and perceived pain intensity: relations to cortisol concentrations after electrocutaneous stimulation in healthy young men. Biopsychosoc Med. 2011;5(1):8.

18. Fries E, Hesse J, Hellhammer J, Hellhammer DH. A new view on hypocortisolism. Psychoneuroendocrinology. 2005 Nov;30(10):1010–6.

19. Rohleder N. Stress and inflammation – The need to address the gap in the transition between acute and chronic stress effects. Psychoneuroendocrinology. 2019 Jul;105:164–71.

20. Yang N, Ray DW, Matthews LC. Current concepts in glucocorticoid resistance. Steroids. 2012 Sep;77(11):1041–9.

21. Szabo YZ, Slavish DC, Graham-Engeland JE. The effect of acute stress on salivary markers of inflammation: A systematic review and meta-analysis. Brain Behav Immun. 2020 Aug;88:887–900.

22. Giletta M, Slavich GM, Rudolph KD, Hastings PD, Nock MK, Prinstein MJ. Peer victimization predicts heightened inflammatory reactivity to social stress in cognitively vulnerable adolescents. Journal of Child Psychology and Psychiatry. 2018 Feb;59(2):129–39.

23. Saban KL, Mathews HL, Bryant FB, Tell D, Joyce C, DeVon HA, et al. Perceived discrimination is associated with the inflammatory response to acute laboratory stress in women at risk for cardiovascular disease. Brain Behav Immun. 2018 Oct;73:625–32.

24. Slavish DC, Graham-Engeland JE, Smyth JM, Engeland CG. Salivary markers of inflammation in response to acute stress. Brain Behav Immun. 2015 Feb;44:253–69.

25. Chojnowska S, Ptaszyńska-Sarosiek I, Kępka A, Knaś M, Waszkiewicz N. Salivary Biomarkers of Stress, Anxiety and Depression. J Clin Med. 2021 Feb 1;10(3):517.

26. La Fratta I, Tatangelo R, Campagna G, Rizzuto A, Franceschelli S, Ferrone A, et al. The plasmatic and salivary levels of IL-1β, IL-18 and IL-6 are associated to emotional difference during stress in young male. Sci Rep. 2018 Dec 14;8(1):3031.

27. Auer BJ, Calvi JL, Jordan NM, Schrader D, Byrd-Craven J. Communication and social interaction anxiety enhance interleukin-1 beta and cortisol reactivity during high-stakes public speaking. Psychoneuroendocrinology. 2018 Aug;94:83–90.

28. Miller GE, Cohen S, Ritchey AK. Chronic psychological stress and the regulation of pro-inflammatory cytokines: A glucocorticoid-resistance model. Health Psychology. 2002;21(6):531–41.

29. Dickerson SS, Kemeny ME. Acute Stressors and Cortisol Responses: A Theoretical Integration and Synthesis of Laboratory Research. Psychol Bull. 2004;130(3):355–91.

30. Yaribeygi H, Panahi Y, Sahraei H, Johnston TP, Sahebkar A. The impact of stress on body function: A review. EXCLI J. 2017;16:1057–72.

31. Wirtz PH, von Känel R. Psychological Stress, Inflammation, and Coronary Heart Disease. Curr Cardiol Rep. 2017 Nov 20;19(11):111.

32. Onyango AN. Cellular Stresses and Stress Responses in the Pathogenesis of Insulin Resistance. Oxid Med Cell Longev. 2018 Jul 9;2018:1–27.

33. Dai S, Mo Y, Wang Y, Xiang B, Liao Q, Zhou M, et al. Chronic Stress Promotes Cancer Development. Front Oncol. 2020 Aug 19;10.

34. Gao H, Wang X, Huang M, Qi M. Chronic academic stress facilitates response inhibition: Behavioral and electrophysiological evidence. Cogn Affect Behav Neurosci. 2022 Jun 15;22(3):533–41.

35. Manolova MS, Stefanova VP, Manchorova-Veleva NA, Panayotov I V., Levallois B, Tramini P, et al. A Five-year Comparative Study of Perceived Stress Among Dental Students at Two European Faculties. Folia Med (Plovdiv). 2019 Mar 1;61(1):134–42.

36. Knowles SR, Nelson EA, Palombo EA. Investigating the role of perceived stress on bacterial flora activity and salivary cortisol secretion: A possible mechanism underlying susceptibility to illness. Biol Psychol. 2008 Feb;77(2):132–7.

37. Frazier P, Gabriel A, Merians A, Lust K. Understanding stress as an impediment to academic performance. Journal of American College Health. 2019 Aug 18;67(6):562–70.

38. Chyu EPY, Chen JK. The Correlates of Academic Stress in Hong Kong. Int J Environ Res Public Health. 2022 Mar 28;19(7):4009.

39. S. MF,. JJ,. ME. Estrés académico en estudiantes universitarios. Investig Cienc [Internet]. 2020;28:75–83. Available from: https://www.redalyc.org/articulo.oa?id=67462875008

40. Jiménez-Mijangos LP, Rodríguez-Arce J, Martínez-Méndez R, Reyes-Lagos JJ. Advances and challenges in the detection of academic stress and anxiety in the classroom: A literature review and recommendations. Educ Inf Technol (Dordr). 2023 Apr 28;28(4):3637–66.

41. Barraza A. Inventario SISCO estrés académico. Propiedades psicométricas. Revista PsicologiaCientifica.com. 2007 Feb;9(13).

42. Barraza Macías A. INVENTARIO SISCO SV-21. Inventario SIStémico COgnoscitivista para el estudio del estrés académico. Segunda versión de 21 ítems. 2018.

43. Castillo AG, Saez K, Perez C, Castillo Navarrete JL. Validity and reliability of SISCO inventory of academic stress among health students in Chile. J Pak Med Assoc. 2018 Dec;68(12):1759–62.

44. Guzmán-Castillo A, Bustos N. C, Zavala S. W, Castillo-Navarrete JL. Inventario SISCO del estrés académico: revisión de sus propiedades psicométricas en estudiantes universitarios. Terapia psicológica. 2022 Jul;40(2):197–211.

45. Castillo-Navarrete JL, Bustos C, Guzman-Castillo A, Zavala W. Academic stress in college students: descriptive analyses and scoring of the SISCO-II inventory. PeerJ. 2024 Mar 12;12:e16980.

46. Messner B, Bernhard D. Smoking and Cardiovascular Disease. Arterioscler Thromb Vasc Biol. 2014 Mar;34(3):509–15.

47. Morris NL, Ippolito JA, Curtis BJ, Chen MM, Friedman SL, Hines IN, et al. Alcohol and inflammatory responses: Summary of the 2013 Alcohol and Immunology Research Interest Group (AIRIG) meeting. Alcohol. 2015 Feb;49(1):1–6.

48. Green ES, Arck PC. Pathogenesis of preterm birth: bidirectional inflammation in mother and fetus. Semin Immunopathol. 2020 Aug 7;42(4):413–29.

49. Castillo-Navarrete J, Guzmán-Castillo A, Bustos N. C, Zavala S. W, Vicente P. B. Propiedades Psicométricas del Inventario SISCO-II de Estrés Académico. Revista Iberoamericana de Diagnóstico y Evaluación – e Avaliação Psicológica. 2020 Jul;56(3):101.

50. Ponce P, del Arco A, Loprinzi P. Physical Activity versus Psychological Stress: Effects on Salivary Cortisol and Working Memory Performance. Medicina (B Aires). 2019 Apr 30;55(5):119.

51. Fiksdal A, Hanlin L, Kuras Y, Gianferante D, Chen X, Thoma M V., et al. Associations between symptoms of depression and anxiety and cortisol responses to and recovery from acute stress. Psychoneuroendocrinology. 2019 Apr;102:44–52.

52. Zefferino R, Fortunato F, Arsa A, Di Gioia S, Tomei G, Conese M. Assessment of Stress Salivary Markers, Perceived Stress, and Shift Work in a Cohort of Fishermen: A Preliminary Work. Int J Environ Res Public Health. 2022 Jan 8;19(2):699.

53. Pulopulos MM, Baeken C, De Raedt R. Cortisol response to stress: The role of expectancy and anticipatory stress regulation. Horm Behav. 2020 Jan;117:104587.

54. de Punder K, Heim C, Entringer S. Association between chronotype and body mass index: The role of C-reactive protein and the cortisol response to stress. Psychoneuroendocrinology. 2019 Nov;109:104388.

55. Smyth N, Hucklebridge F, Thorn L, Evans P, Clow A. Salivary Cortisol as a Biomarker in Social Science Research. Soc Personal Psychol Compass. 2013 Sep 2;7(9):605–25.

56. Kudielka BM, Hellhammer DH, Wüst S. Why do we respond so differently? Reviewing determinants of human salivary cortisol responses to challenge. Psychoneuroendocrinology. 2009 Jan;34(1):2–18.

57. Palumbo ML, Prochnik A, Wald MR, Genaro AM. Chronic Stress and Glucocorticoid Receptor Resistance in Asthma. Clin Ther. 2020 Jun;42(6):993–1006.

58. Reschke-Hernández AE, Okerstrom KL, Bowles Edwards A, Tranel D. Sex and stress: Men and women show different cortisol responses to psychological stress induced by the Trier social stress test and the Iowa singing social stress test. J Neurosci Res. 2017 Jan 2;95(1–2):106–14.

59. Nijakowski K, Gruszczyński D, łaganowski K, Furmańczak J, Brożek A, Nowicki M, et al. Salivary Morning Cortisol as a Potential Predictor for High Academic Stress Level in Dental Students: A Preliminary Study. Int J Environ Res Public Health. 2022 Mar 7;19(5):3132.

60. Helbig S, Backhaus J. “Sex differences in a real academic stressor, cognitive appraisal and the cortisol response.” Physiol Behav. 2017 Oct;179:67–74.

61. Steptoe A, Hamer M, Chida Y. The effects of acute psychological stress on circulating inflammatory factors in humans: A review and meta-analysis. Brain Behav Immun. 2007 Oct;21(7):901–12.

62. Newton TL, Fernandez-Botran R, Lyle KB, Szabo YZ, Miller JJ, Warnecke AJ. Salivary cytokine response in the aftermath of stress: An emotion regulation perspective. Emotion. 2017 Sep;17(6):1007–20.

63. Ballestar-Tarín ML, Ibáñez-del Valle V, Mafla-España MA, Cauli O, Navarro-Martínez R. Increased Salivary IL-1 Beta Level Is Associated with Poor Sleep Quality in University Students. Diseases. 2023 Oct 5;11(4):136.

64. Paudel D, Morikawa T, Yoshida K, Uehara O, Giri S, Neopane P, et al. Chronic stress-induced elevation of IL-1β in the saliva and submandibular glands of mice. Med Mol Morphol. 2020 Dec 6;53(4):238–43.

65. Goetz SM, Lucas T. C-reactive protein in saliva and dried blood spot as markers of stress reactivity in healthy African–Americans. Biomark Med. 2020 Apr;14(5):371–80.

66. Seizer L, Stasielowicz L, Löchner J. Timing matters: A meta-analysis on the dynamic effect of stress on salivary immunoglobulin. Brain Behav Immun. 2024 Jul;119:734–40.

67. Bosch JA, Ring C, de Geus EJC, Veerman ECI, Nieuw Amerongen A V. Stress and secretory immunity. In 2002. p. 213–53.

68. Ding L, Chen X, Cheng H, Zhang T, Li Z. Advances in IgA glycosylation and its correlation with diseases. Front Chem. 2022 Sep 27;10.

69. Al-Rabiaah A, Temsah MH, Al-Eyadhy AA, Hasan GM, Al-Zamil F, Al-Subaie S, et al. Middle East Respiratory Syndrome-Corona Virus (MERS-CoV) associated stress among medical students at a university teaching hospital in Saudi Arabia. J Infect Public Health. 2020 May;13(5):687–91.

70. Dopelt K, Houminer-Klepar N. War-Related Stress among Israeli College Students Following 7 October 2023 Terror Attack in Israel. Eur J Investig Health Psychol Educ. 2024 Jul 30;14(8):2175–86.

71. Sood P. Psychological stressors as interventions: Good out of the evil. Frontiers in Bioscience. 2012;S4(1):43.

72. Love MF, Sharrief A, Chaoul A, Savitz S, Beauchamp JES. Mind-Body Interventions, Psychological Stressors, and Quality of Life in Stroke Survivors. Stroke. 2019 Feb;50(2):434–40.

73. Biltz RG, Sawicki CM, Sheridan JF, Godbout JP. The neuroimmunology of social-stress-induced sensitization. Nat Immunol. 2022 Nov 11;23(11):1527–35.

