## Supplementary figures and images for "Academic stress through salivary biomarkers: A multivariate analysis of cortisol, IL-1β, CRP, and IgA levels and their variations as a function of sex"

### Suplemental file correlation CRP IL-1B men.docx

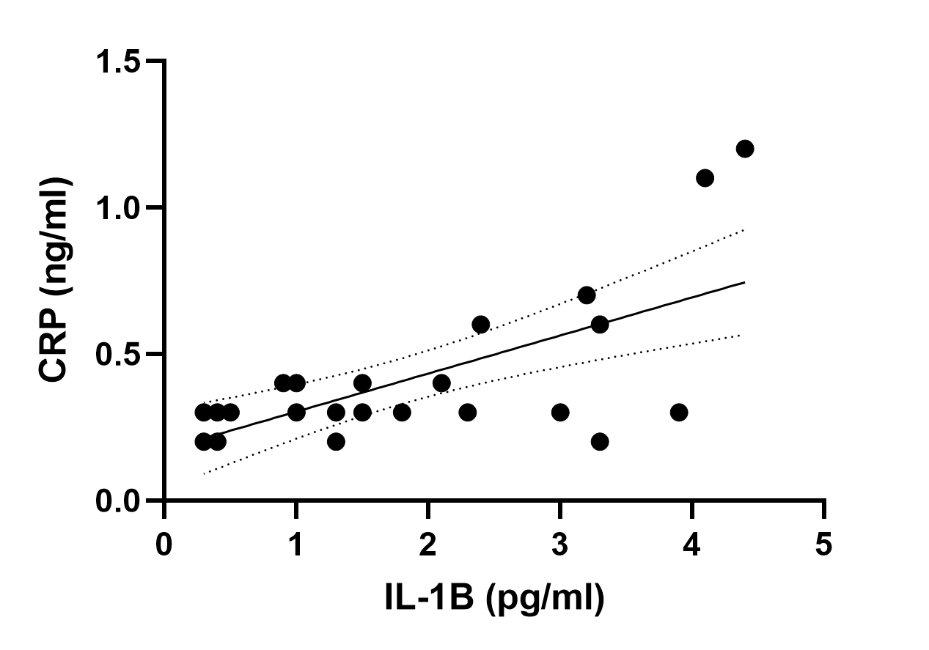
